# The hedgehog signaling pathway is expressed in the adult mouse hypothalamus and modulated by fasting

**DOI:** 10.1101/2021.06.09.447747

**Authors:** Patrick J. Antonellis, Staci E. Engle, Kathryn M. Brewer, Nicolas F. Berbari

## Abstract

The importance of the primary cilium was initially highlighted by the class of human genetic disorders known as ciliopathies. Patients with ciliopathies such as Bardet-Biedl and Alström syndrome exhibit hyperphagia-associated obesity as a core clinical phenotype. How primary cilia contribute to energy homeostasis and feeding behavior is complex and unclear, but cilia appear important in both developmental and homeostatic processes. Primary cilia are important signaling centers, required for hedgehog signaling and localization of specific G protein-coupled receptors (GPCRs) with known roles in feeding behavior in mammals. The hedgehog pathway is best known for its role in developmental patterning, but now has recognized roles in adult tissues as well. In the postnatal brain, cilia and hedgehog signaling are important for growth and maintenance of neural progenitors, however, the role of hedgehog signaling in the differentiated adult brain is less clear. Here, we provide a detailed analysis of the expression of core components of the hedgehog signaling pathway in the adult mouse hypothalamus with an emphasis on feeding centers. We show that hedgehog pathway genes continue to be expressed in differentiated neurons important for regulation of feeding behavior. Furthermore, we demonstrate for the first time that pathway activity is regulated at the transcriptional level by fasting. These data suggest that hedgehog signaling is involved in the proper functioning of brain regions which regulate feeding behavior and that hedgehog pathway dysfunction may play a role in the obesity observed in certain ciliopathies.

**Significance Statement:** Here we investigate the expression of hedgehog pathway components in the adult mouse hypothalamus. Using dual labeling in situ hybridization we show that core components of the signaling pathway are expressed in multiple neuronal cell types within the hypothalamic feeding centers. Our findings also support previous findings that astrocytes are responsive to hedgehog signaling, as determined by *Gli1* and *Ptch1* expression. Using qPCR analysis, we show that hypothalamic hedgehog pathway activity is upregulated in response to fasting and this response is nuclei specific. These data not only provide a more detailed understanding of hedgehog pathway expression in the adult mouse hypothalamus but also provide direct evidence of a novel role for hedgehog signaling in the physiological response to fasting.

## 1 Introduction

Primary cilia are found on almost all mammalian cell types and their dysfunction underlies a class of human genetic disorders known as ciliopathies (Reiter and Leroux, 2017). Obesity is a core clinical phenotype of certain ciliopathies such as Bardet-Biedl (BBS, OMIM #209900) and Alström (ALMS, OMIM #203800) syndromes (Marshall et al., 2011; Forsythe et al., 2018). Ciliopathy patients present a wide range of clinical symptoms, some of which are also associated with genetic defects in hedgehog signaling such as bone, limb patterning and genitalia malformations (Umehara et al., 2000; Gao et al., 2001; Hellemans et al., 2003; Mujahid et al., 2018; Khan et al., 2019). Furthermore, the ciliopathy Carpenter syndrome (OMIM #20100) results from mutations in Rab23, a negative regulator of hedgehog signaling. Clinical features of patients with Carpenter syndrome include both skeletal malformations, and obesity (Jenkins et al., 2007; Alessandri et al., 2010). Thus, these genetic disorders implicate a potential role for hedgehog signaling in regulation of energy homeostasis in humans.

Ciliopathy mouse models, as well as conditional animal models of cilia loss, have implicated hypothalamic neuronal cilia-mediated feeding behaviors in obesity (Davenport et al., 2007; Siljee et al., 2018; Wang et al., 2019; Rouabhi et al., 2021; Sun et al., 2021). Neuronal primary cilia preferentially localize G protein-coupled receptors (GPCRs) known to be important for regulation of energy homeostasis and feeding behavior, such as neuropeptide Y receptors 2 and 5, melanin-concentrating hormone receptor 1 (MCHR1) and melanocortin-4 receptor (MC4R) (Berbari et al., 2008; Loktev and Jackson, 2013; Siljee et al., 2018). However, the impact of ciliary localization on their signaling capabilities is not well understood. We have previously demonstrated in primary mouse hypothalamic neurons *in vitro*, interactions between a ciliary GPCR and hedgehog signaling, suggesting that the hedgehog pathway may modulate GPCR activity at the cilium in differentiated neurons (Bansal et al., 2019).

In mammals, the hedgehog pathway is dependent on the primary cilium as an organizing center (Huangfu et al., 2003; Tran et al., 2008; Willaredt et al., 2008; Gorivodsky et al., 2009; Stottmann et al., 2009). Components of the hedgehog pathway such as patched (PTCH1), smoothened (SMO) and the Glioma (Gli) transcription factors GLI2 and GLI3 dynamically localize to the primary cilia (Corbit et al., 2005; Haycraft et al., 2005; Rohatgi et al., 2007). Upon binding to the ligand sonic hedgehog (SHH), PTCH1 leaves the cilia allowing for SMO to enter primary cilia. GLI2 is activated into its transcriptional activator (GLI2A) form while GLI3R formation is inhibited, leading to expression of Gli target genes, which includes *Ptch1* and *Gli1*, (Chen and Struhl, 1996; Marigo and Tabin, 1996; Bai et al., 2004; Fuccillo et al., 2006). Numerous other genes are known to regulate hedgehog signaling in embryonic development, such as *Gpr161*, an orphan GPCR shown to localize to cilia and negatively regulate pathway activity (Mukhopadhyay et al., 2013).

Beyond embryonic development, cilia and hedgehog signaling continue to play an important role in the postnatal brain for growth and the maintenance of neural progenitors (Machold et al., 2003; Chizhikov et al., 2007; Breunig et al., 2008; Han et al., 2008; Vaillant and Monard, 2009). However, the role of hedgehog signaling in the adult hypothalamus is less clear. *In situ* hybridization studies have shown that *Shh*, *Ptch1*, and *Smo* mRNA are expressed in several regions of adult rat brain, including the hypothalamus (Traiffort et al., 1998; Traiffort et al., 1999; Traiffort et al., 2001; Banerjee et al., 2005). Here we characterize in greater detail the expression and transcriptional activity of the hedgehog pathway in the feeding centers of the adult hypothalamus *in vivo*. We also demonstrate that hedgehog pathway activity changes based on feeding status and this response is absent following the onset of obesity, suggesting a role for hedgehog signaling in the modulation of adult behaviors.

## 2 Materials and Methods

### 2.1 Animals

All procedures were approved by the Institutional Animal Care and Use Committee at Indiana University-Purdue University Indianapolis. Male C57BL/6J (stock 000664) mice were ordered from The Jackson Laboratory and housed on a standard 12-hour light dark cycle and given food and water *ad libitum* except for experiments as described. Chow fed mice were maintained on a standard chow diet consisting of 13% fat, 67% carbohydrate, and 20% protein caloric content (2014; Envigo). High fat diet (HFD) fed animals were maintained on a diet consisting of 40% fat, 39% carbohydrate, and 21% protein caloric content starting at 6 weeks of age (TD95217; Envigo).

### 2.2 In situ hybridization

Brains from 8 to 10 week old male C57BL/6J mice were harvested and fixed as previously described (Engle et al., 2018). Sections were cut at a thickness of 15 μm and mounted directly on slides then post-fixed with 4% paraformaldehyde for 16 hr at 4LC. Detection of transcripts in brain sections was performed using the RNAscope 2.5 HD Duplex Assay (ACD, Newark, CA). Tissue pretreatment was performed according to technical note 320534 Rev A. Probe hybridization, counterstaining, and mounting of slides was performed according to user manual no. 322500-USM Rev A. Slides were assayed using probes to SHH (Cat No. 314361), SMO (Cat No. 318411), GLI1 (Cat No. 311001), PTCH1 (Cat No. 402811), GPR161 (Cat No. 318111), AGRP (Cat No. 400711-C2), POMC (Cat No. 314081-C2), MC4R (Cat No. 319181-C2), MCHR1 (Cat No. 317491-C2), or GFAP (Cat No. 313211-C2) transcripts (ACD). Sections were counterstained with hematoxylin, dehydrated, and mounted using VectaMount (Vectorlabs, Burlingame, CA). Slides with positive control probe (PPIB-C1/POLR2A-C2; Cat No. 321651) and negative control probe (DapB; Cat no. 320751) were ran with each experiment. At least 3 animals were analyzed for each group.

### 2.3 Quantitative Real-Time PCR

RNA was isolated, cDNA prepared, and quantitative real-time PCR performed as previously described (Bansal et al., 2019). Assays-on-Demand Gene expression probes (Applied Biosystems) were as follows: Shh Mm00436528_m1; Ptch1 Mm00436026_m1; Smo Mm01162710_m1; Gli1 Mm00494654_m1; Gpr161: Mm01291057_m1. CT values were normalized to β-actin, and relative expression was calculated by the ΔΔCT method and fold change was calculated by normalizing relative expression to the proper control.

### 2.4 Experimental Design and Statistical Analyses

Whole hypothalamus was collected from 35 to 36 week old, C57Bl/6, lean and obese animals that were either allowed *ad libitum* access to food or fasted overnight. Region specific micropunches were collected from 7 to 8 week old, lean, C57Bl/6 mice using 1.0 mm Militex Biopsy Punch (Electron Microscopy Sciences). There was a minimum of 6 animals per treatment group. Statistical analysis was performed using two-way ANOVA and corrected for multiple comparisons. Differences were considered significant when P < 0.05. Data are presented as mean ± SEM.

## 3 Results

To determine whether *Shh* and components of the signaling pathway are expressed in the adult mouse hypothalamus we performed *in situ* hybridization studies. Using a dual labeling approach, we first assessed whether neurons of one of the feeding centers, the arcuate nucleus of the hypothalamus (ARC), express hedgehog pathway genes. Two major neuronal subtypes within the ARC, the anorexigenic proopiomelanocortin (POMC) expressing neurons and orexigenic agouti-related protein (AgRP)/neuropeptide Y (NPY) coexpressing neurons, are crucial for normal energy homeostasis (Belgardt et al., 2009). Hypothalamic sections from adult, C57/Bl6 mice were labeled with probes to *Shh*, *Ptch1*, *Smo*, *Gli1*, or *Gpr161* and colabeled with probes to either *Pomc* or *Agrp*. In the ARC, we found that all hedgehog pathway genes assayed were coexpressed in neurons expressing *Pomc* (**Figure 1A-D**) and *Agrp* (**Figure 1F-I**). We also found that *Gpr161* is expressed at a relatively high level throughout the ARC (**Figure 1E** and **1J**). For each experiment, sections were labeled with positive and negative control probes (**Supplemental Figure 1**).

**Figure 1:**
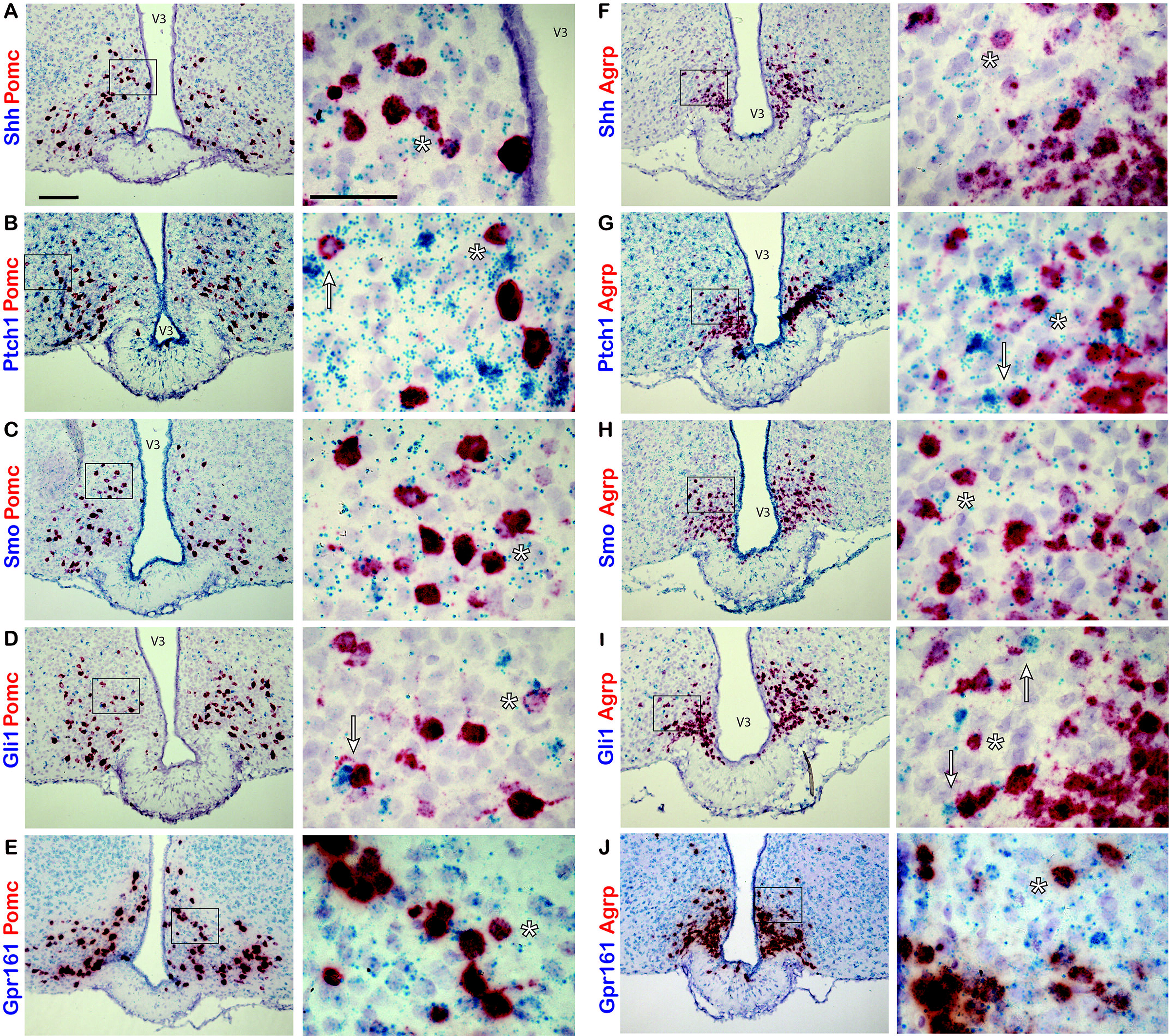
*Hedgehog pathway expression in the adult mouse ARC*. Dual probe *in situ* hybridization labeling of the ARC with probes to hedgehog pathway and neuronal gene transcripts. *Pomc* (**A-E**) and *Agrp* (**F-J**) probes are labeled in red while *Shh* (**A** and **F**), *Ptch1* (**B** and **G**), *Smo* (**C** and **H**), *Gli1* (**D** and **I**), and *Gpr161* (**E** and **J**) probes are labeled in blue. Examples of cells colabeled by both probes are denoted by an asterisk (**\***). *Pomc* or *Agrp* expressing cells adjacent to highly expressing *Ptch1* (**B** and **G**) or *Gli1* (**D** and **I**) cells are denoted by an arrow. Right hand panels are magnified images of region in black box on left hand side. Scale bar 100 μm on left panels and 50 μm right panels. V3 indicates third ventricle.

We next assessed whether *Shh* and its pathway components are also expressed in another feeding center, the paraventricular nucleus of the hypothalamus (PVN). We labeled sections of hypothalamus with probes to *Shh*, *Ptch1*, *Smo*, *Gli1*, or *Gpr161* and colabeled with probes to either *Mchr1* or *Mc4r*. Both MCHR1 and MC4R are known to be important for regulation of energy homeostasis and feeding behavior and localize to primary cilia in neurons (Berbari et al., 2008;Siljee et al., 2018). In the PVN, we observed relatively few *Mc4r* positive neurons with sparse incidence of colabeling with probes to hedgehog pathway genes (**Figure 2A-D**). However, *Mchr1* positive neurons were much more abundant and were frequently colabeled with probes to all hedgehog pathway genes used (**Figure 2F-I**). As in the ARC, *Gpr161* is expressed abundantly throughout the PVN (**Figure 2E** and **J**).

**Figure 2:**
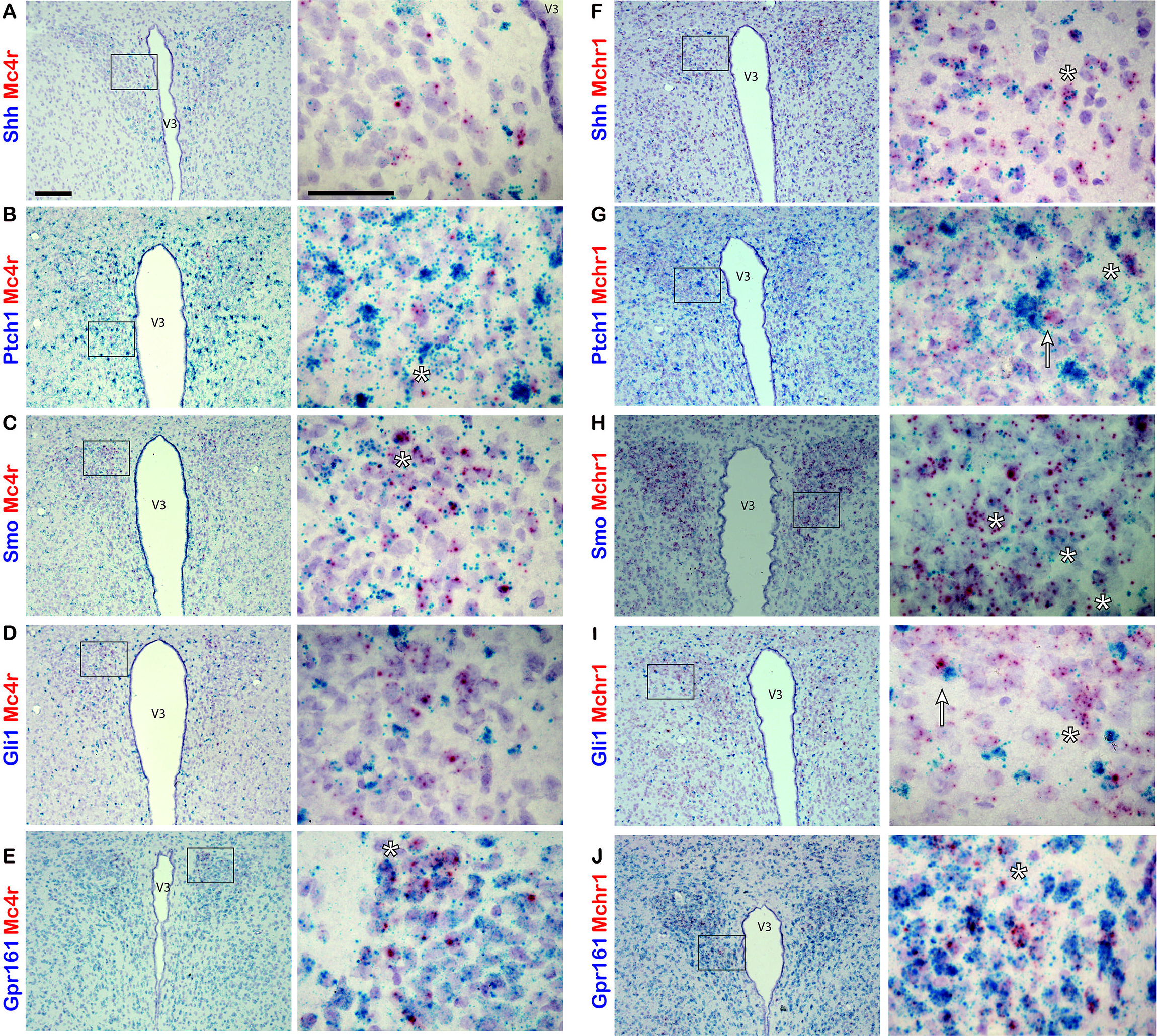
*Hedgehog pathway expression in adult mouse PVN.* Dual probe *in situ* hybridization of probes for hedgehog pathway gene transcripts colabeled with probes to neuronal gene transcripts in the PVN of adult mice. *Mc4r* (**A**-**E**) and *Mchr1* (**F**-**J**) probes are labeled in red while *Shh* (**A** and **F**), *Ptch1* (**B** and **G**), *Smo* (**C** and **H**), *Gli1* (**D** and **I**), and *Gpr161* (**E** and **J**) probes are labeled in blue. Example of cells colabeled by both probes are denoted by an asterisk (**\***). *Mchr1* expressing cells adjacent to highly expressing *Gli1* cells are denoted by an arrow (**I**). Right hand panels are magnified images of region in black box on left hand side. Scale bar 100 μm left panels and 50 μm right panels. V3 indicates third ventricle.

Interestingly, we observed cells highly positive for either *Gli1* or *Ptch1* expression throughout the ARC and PVN. Some of these cells appeared adjacent to neurons expressing *Pomc*, *Agrp*, or *Mchr1* (**Figure 1B, 1D, 1G, 1I**, and **2I**, arrows). Previously it has been reported that astrocytes of the adult mouse brain are responsive to hedgehog signaling and increase *Gli1* expression upon pathway activation (Garcia et al., 2010). Therefore, we sought to determine if these hedgehog responsive cells were astrocytes by colabeling with probes to either *Gli1* or *Ptch1* and glial fibrillary acidic protein (*Gfap*), an astrocyte marker. In the ARC (**Figure 3A** and **C**) and PVN (**Figure 3B** and **D**), cells highly expressing *Gfap* were observed colabeled with *Gli1* or *Ptch1*, suggesting that some of the hedgehog responsive cells adjacent to neurons may be astrocytes in the hypothalamus.

**Figure 3:**
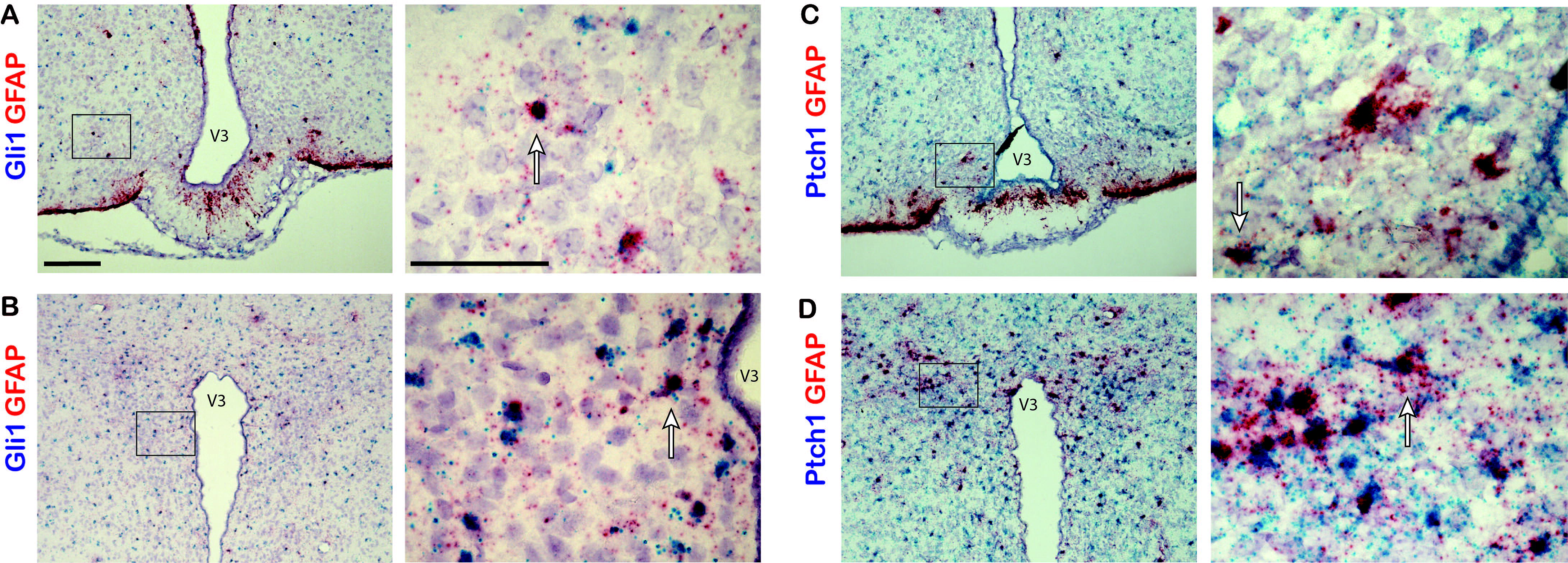
*Gli1 and Ptch1 expression in astrocytes in adult mouse hypothalamus.* Dual probe *in situ* hybridization was performed with probes for *Gli1* or *Ptch1* and astrocyte marker *Gfap* gene transcripts. Sections of the ARC (**A and C**) and PVN (**B and D**) from adult mice were colabeled with probes to *Gli1* or *Ptch1* labeled in blue and probes to *Gfap* labeled in red. Example cells which are highly positive for both probes are denoted by an arrow. Right hand panels are magnified images of region in black box on left hand side.100 μm scale bar for left side panels, 50 μm scale bar on right side. V3 indicates third ventricle.

Since we determined that genes of the hedgehog pathway are expressed in neurons of the adult hypothalamus, we next wanted to determine whether these genes are transcriptionally regulated by physiological changes associated with the normal function of this brain region, such as nutritional status. This was accomplished by gene expression analysis of the whole hypothalamus from chow fed lean and high fat diet (HFD) fed obese animals in the fed and fasted state. HFD fed animals weighed significantly more than their chow fed counterparts at 35 weeks of age (57.1g +/− 0.97 versus 34.8g +/− 0.75; mean +/− SEM; p < 0.05, Student’s t-test). Lean and obese animals were allowed either *ad libitum* access to food or fasted overnight, then whole hypothalamic RNA was collected for qPCR analysis. In lean animals there was a significant increase in both *Shh* and *Gli1* expression in the hypothalamus after an overnight fast (**Figure 4A**). The expression of *Ptch1* also was elevated in fasted compared to fed animals, but this effect was not significant (**Figure 4A**). Strikingly, there were no significant changes in gene expression in the hypothalamus of obese fasted vs fed animals (**Figure 4A**). Since transcriptional regulation was observed in response to fasting in control chow fed animals at the level of the whole hypothalamus, we wanted to assess which specific nuclei were contributing to this effect. Once again control, normal weight C57Bl/6J animals were either allowed *ad libitum* access to food or fasted overnight. Micropunches were then collected from the cortex, ventromedial hypothalamus (VMH), PVN, and ARC for qPCR analysis. We found the fasting induced increase in *Gli1* expression in the whole hypothalamus is driven by increases in *Gli1* expression specifically in the VMH and PVN but not the ARC (**Figure 4B**). We also found that *Gli1* was upregulated, to a lesser extent, in the cortex (**Figure 4B**). Additionally, expression of *Smo* was increased in the VMH, while *Shh* gene expression was reduced in the ARC following an overnight fast (**Figure 4B**). Finally, we performed qPCR on whole hypothalamus and cortex, a brain region known to exhibit hedgehog pathway activity (Garcia et al., 2010; Harwell et al., 2012; Hill et al., 2019), collected from adult animals. We found that all hedgehog pathway genes measured were more highly expressed in the hypothalamus than the cortex (**Supplemental Figure 2).** Overall, these data demonstrate that not only does expression of the hedgehog pathway continue into adulthood in the hypothalamus, a region critical for energy homeostasis, but that specific nuclei respond with transcriptional changes based upon feeding status.

**Figure 4:**
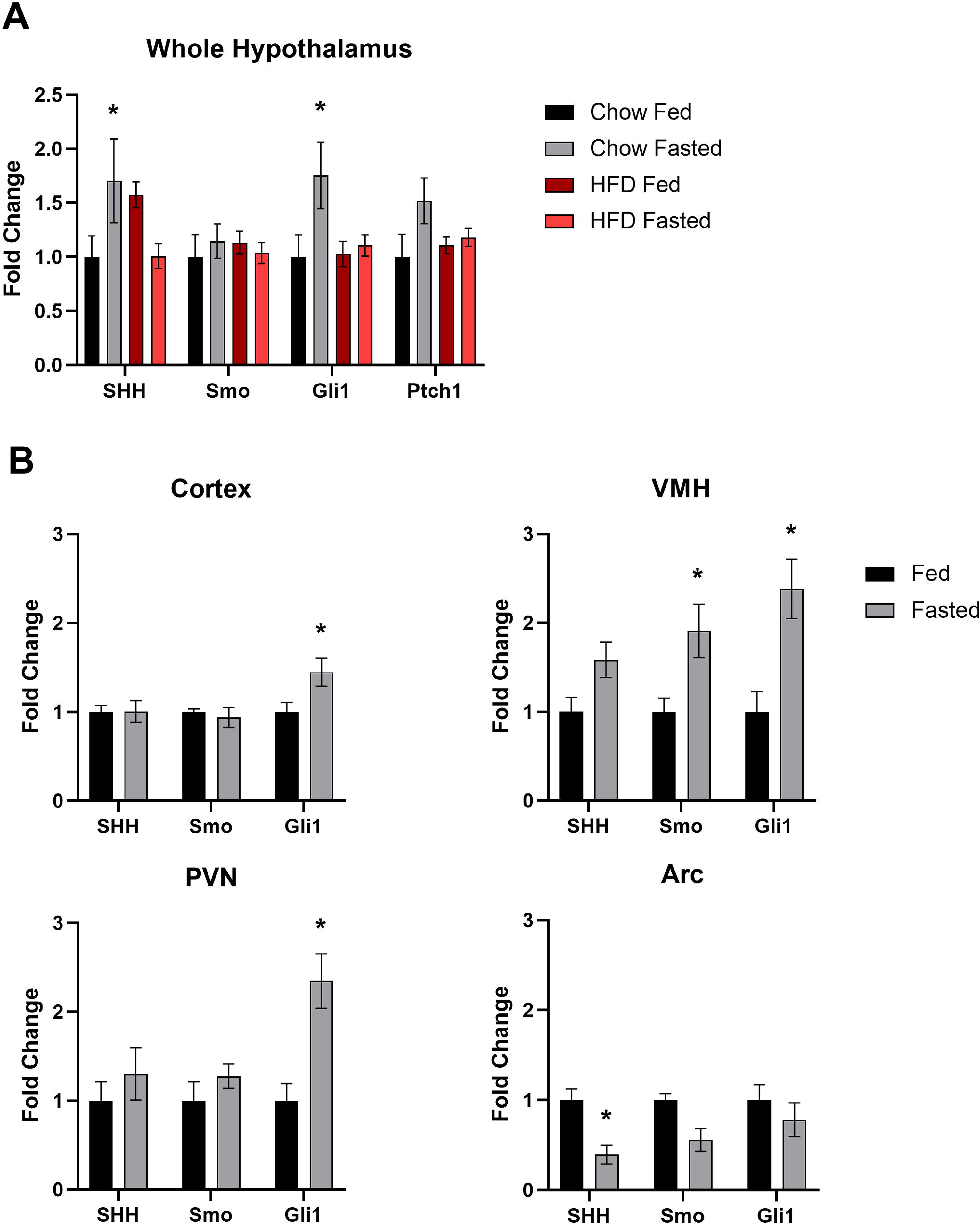
*Feeding status mediated changes in hypothalamic hedgehog pathway gene expression.* (**A**) Hedgehog pathway gene expression in the whole hypothalamus of adult mice. Lean animals fed a standard chow or obese animals fed high-fat diet were allowed *ad libitum* access to food or fasted overnight. A minimum of 6 animals were used per treatment group. Whole hypothalamic RNA was used for qPCR. (**B**) Hedgehog pathway gene expression in brain micropunches of adult mice. A total of 16 animals, 8 per treatment group, fed a chow diet were allowed *ad libitum* access to food or fasted overnight. Micropunches were taken from specific nuclei of the hypothalamus and cortex for qPCR analysis. Asterisks denote a significance value of p < 0.05.

## 4 Discussion

Primary cilia are crucial for mediating hedgehog signaling in mammals and furthermore, certain ciliopathies are associated with pediatric obesity (Engle et al., 2021). Therefore, we focused our efforts on evaluating expression of the hedgehog pathway in hypothalamic feeding centers of adult mice. Because reliable commercial antibodies for many pathway proteins are currently unavailable, making immunolabelling studies difficult, we utilized a dual-labeling *in situ* hybridization approach which allowed for sensitive detection of hedgehog pathway gene transcripts in specific adult neurons. Our *in situ* data revealed the broad expression of *Shh* and core pathway components throughout the hypothalamus of adult mice. We found that mRNA for *Shh*, *Ptch1*, *Smo*, *Gli1*, and *Gpr161* could be detected in both the ARC and PVN. Within the ARC, each one of these transcripts are detected in neurons co-expressing *Pomc* or *Agrp*. While a similar labeling pattern was observed in the PVN, transcripts for hedgehog pathway genes were more readily found colabeled with probes to *Mchr1* than *Mc4r*. However, this could potentially be due to the relatively low abundance of MC4R expressing neurons or low mRNA expression for this particular GPCR. Our results expand upon prior studies which found hedgehog pathway expression in the adult rat brain (Traiffort et al., 1998; Traiffort et al., 1999; Banerjee et al., 2005) by demonstrating that neurons in two nuclei of the hypothalamus important for regulation of feeding behavior express *Shh* and members of its signaling pathway. Somewhat surprisingly, we also found that *Gpr161* is expressed abundantly throughout both the ARC and PVN. This is contrary to previously published findings which used digoxigenin probes *for in situ* analysis of the adult mouse brain and showed a more restricted expression pattern of *Gpr161* in the nucleus accumbens and amygdala with low hypothalamic expression (Ehrlich et al., 2018). Analysis of the amygdala and accumbens was outside the scope of the present study, and relative expression of *Gpr161* between these brain regions was not determined. However, a thorough understanding of this GPCR negative regulator of hedgehog signaling in the adult brain may reveal themes for its roles in cilia mediated behaviors.

Interestingly, in both the ARC and PVN we observed cells with high expression of either *Ptch1* or *Gli1* immediately adjacent to neurons expressing *Pomc*, *Agrp*, or *Mchr1*. By colabeling sections of hypothalamus with probes to *Gfap* and *Gli1* or *Ptch1 we* were able to identify some of these hedgehog responsive cells as *Gfap* positive astrocytes. While neurons of the adult mouse hypothalamus express *Shh* and components required for its signal transduction, the cells which are most responsive to hedgehog signaling, as indicated by high levels of *Gli1* and *Ptch1* expression, may in fact be astrocytes. This supports previous findings which showed that not only do neurons produce *Shh*, but subpopulations of mature astrocytes in the forebrain are responsive to hedgehog signaling (Garcia et al., 2010). It remains unclear why these *Gfap* positive cells outside of known neurogenic niches exhibit high levels of *Gli1* and *Ptch1* relative to neighboring neurons.

There is growing evidence suggesting that the hedgehog pathway is involved in regulation of whole-body energy homeostasis. It has been demonstrated in both the fat-body of drosophila and adipose tissue of mice that hedgehog signaling regulates adipocyte differentiation (Suh et al., 2006; Pospisilik et al., 2010). Circulating forms of Hedgehog have been detected and during drosophila larval development shown to be secreted from the gut and act on multiple tissues to coordinate development with nutrient availability (Palm et al., 2013; Rodenfels et al., 2014). Given these findings, we also analyzed the expression of *Shh*, *Smo*, and *Gli1* in the adult mouse hypothalamus in response to changes in metabolic state and nutritional status. We compared transcriptional regulation of the hedgehog pathway in both lean, control diet fed animals and obese, HFD fed animals. Animals fed a HFD are a well-established model of non-insulin dependent type II diabetes which exhibit many hallmarks of metabolic dysfunction such as reduced glucose tolerance (Surwit et al., 1988; Fontaine and Davis, 2016). In the whole hypothalamus, expression of *Gli1* was upregulated in lean mice following an overnight fast. Congruent with this finding, expression of *Ptch1* exhibited on non-significant upregulation in lean, fasted animals. In contrast, we observed no significant transcriptional regulation in the hypothalamus of fasted obese animals compared to fed controls. These data suggest that hedgehog signaling is involved in the physiological response to fasting and may be dysregulated in obese animals. To determine if this fasting-induced upregulation of *Gli1* is a response generated by the whole hypothalamus or specific nuclei, micropunches were collected from hypothalamic nuclei as well as the cortex. We found that in fasted animals *Gli1* expression was elevated specifically in the VMH, PVN, and to a lesser extent the cortex, but not the ARC. These data suggest that increased hedgehog pathway activity upon fasting in the hypothalamus is primarily driven by the VMH and PVN. Future studies will determine the cell types responsible within these nuclei for this increase in *Gli1* expression. Additional work is also required to determine the source of pathway activation observed in these studies. It is possible ligand is produced outside of the hypothalamus, therefore, further analysis could potentially reveal the primary source of *Shh* following an overnight fast.

Taken as a whole, this data shows that neurons of the hypothalamus express both *Shh* and members of its signaling pathway required for signal transduction and that activity of this pathway is upregulated in response to fasting in discrete hypothalamic nuclei. Given our *in situ* data identifying astrocytes as being highly positive for *Gli1* and *Ptch1* in the hypothalamus, it is possible that astrocytes or other support cells are primarily responsible for this fasting induced upregulation of hedgehog signaling. We have previously shown in primary hypothalamic cultures consisting of both neurons and glia, that modulation of the hedgehog pathway alters the electrophysiological response to melanin-concentrating hormone (Bansal et al., 2019). Interestingly, it has also been demonstrated in primary cortical cultures that the presence of astrocytes alters the response of neurons to agonism of the hedgehog pathway (Ugbode et al., 2017). Furthermore, astrocyte specific inhibition of hedgehog signaling *in vivo* was shown to disrupt early postnatal organization and remodeling of cortical synapses resulting in increased neuronal excitability (Hill et al., 2019). Together, these findings suggest novel potential roles for hedgehog signaling outside of its roles as a classical developmental morphogen or in stem cell niche regulation.

The data presented here on the expression and transcriptional regulation of the hedgehog pathway in the adult mouse hypothalamus lays the foundation for future mechanistic studies to determine its role in the proper functioning of the hypothalamus. Given that mammalian hedgehog signaling is coordinated by primary cilia, our future studies will focus on how hypothalamic hedgehog expression may contribute to the obesity phenotype seen in ciliopathies such as Bardet-Biedl syndrome and Alström syndrome (Marshall et al., 2011; Forsythe et al., 2018). Interestingly, certain ciliopathy clinical features such as skeletal and external genitalia abnormalities are also observed in patients with genetic defects in the hedgehog pathway (Umehara et al., 2000; Gao et al., 2001; Hellemans et al., 2003; Mujahid et al., 2018; Khan et al., 2019). Therefore, it would be of interest to determine whether hedgehog signaling is dysregulated in the hypothalamus of animal ciliopathy models. Further mechanistic studies are needed to determine whether hedgehog signaling modulates neuronal activity critical for the physiological response to fasting and if genetic modulation of the hedgehog pathway in the hypothalamus alters feeding behavior. In conclusion, elucidating the involvement of this developmentally important signaling pathway in feeding behavior and body composition is an exciting new avenue of the hedgehog pathway to explore. Greater understanding of the hedgehog pathway in adult energy homeostasis may also reveal common themes for this pathway in regulation of other behaviors.

## Acknowledgments

This work was funded by National Institute of Diabetes and Digestive and Kidney Diseases R01 DK114008 to NFB.

---

**Supplemental Figure 1:**
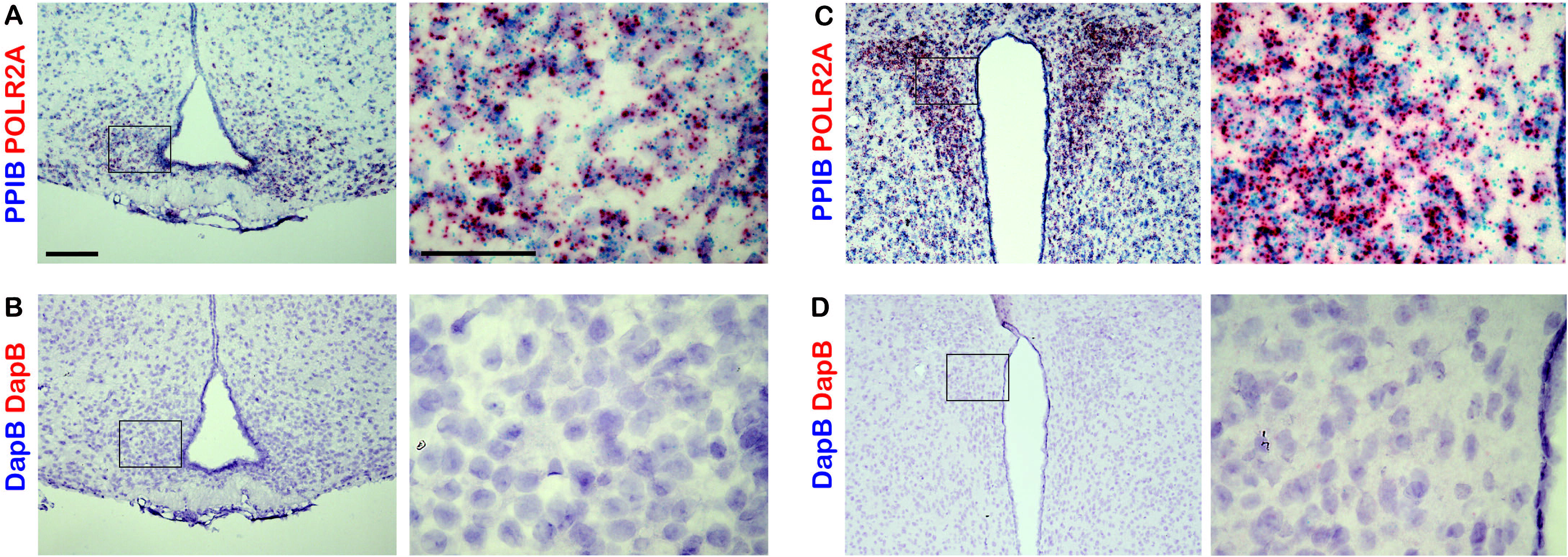
*Controls for dual probe in situ hybridization studies*. Sections of adult mouse ARC and PVN were labeled with either positive control probes targeting the mouse genes *Ppib* and *Polr2a* (**A** and **C**) or negative control probes to the bacterial gene *dapB* (**B** and **D**). Right hand panels are magnified images of region in black box on left hand side. Scale bar 100 μm on left panels and 50 μm right panels

**Supplemental Figure 2:**
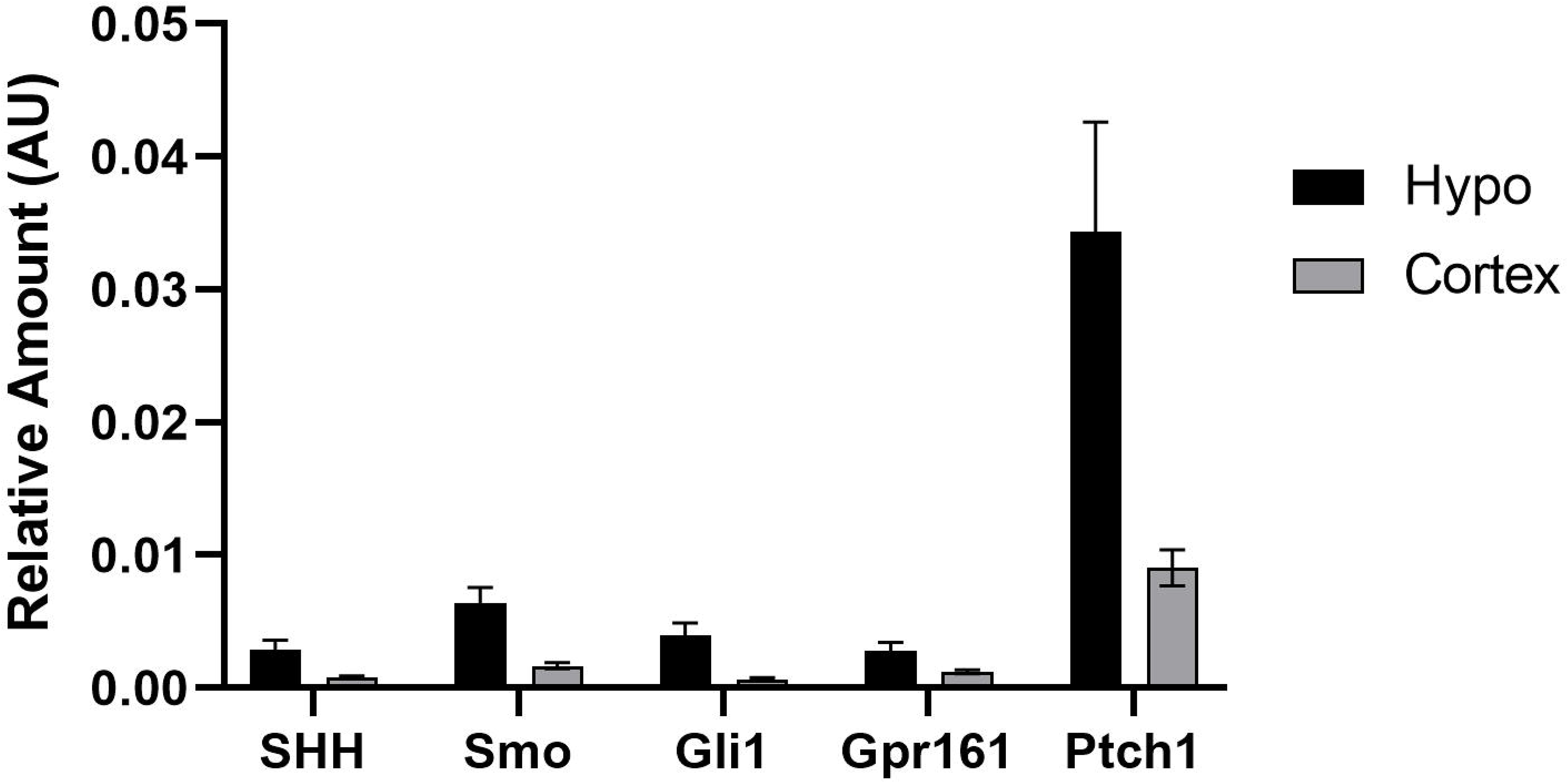
*Expression of selected hedgehog pathway transcripts in the adult mouse brain.* Whole hypothalamus and cortex were collected for qPCR analysis from 6 adult animals allowed *ad libitum* access to food.

